# Amino Acid Metabolic Signature Links Pediatric Environmental Enteric Dysfunction to Maternal-Fetal Growth Restriction Across Two Independent Cohorts

**DOI:** 10.64898/2026.10.25.760166

**Authors:** Shahzaib Khan, Nathaniel Ribeiro, Waqasuddin Khan, Nazanin Moradinasab, Connor Moore, Javairia Khalid, M. Farrukh Qazi, M. Mohsin Mannan, Najeeha Talat Iqbal, Lauren J. Klein, Adam Greene, Asad Ali, Fyezah Jehan, M. Imran Nisar, Sana Syed

**Affiliations:** SUNY Downstate Health Sciences University, Brooklyn, NY; Duke University Medical Center, Durham, NC; Department of Pediatrics and Child Health, Medical College, The Aga Khan University, Karachi-74800, Pakistan; CITRIC Center for Bioinformatics and Computational Biology, Department of Pediatrics and Child Health, Medical College, The Aga Khan University, Karachi-74800, Pakistan; University of Virginia, Charlottesville, VA; Vanderbilt University Medical Center, Nashville, TN; Department of Community Health Sciences, Medical College; Biological and Biomedical Sciences, Aga Khan University, Karachi-74800, Pakistan

**Author notes:** Correspondence: Sana Syed, MD, MSCR, MSDS Division of Pediatric Gastroenterology, Hepatology & Nutrition, Duke University Medical Center, Duke Clinical Research Institute, 300 W Morgan St, Suite 800, Durham, NC 27705, USA. co-first Authors (Shahzaib Khan, Nathaniel Ribeiro). co-last, co-senior, equal contributions (Fyezah Jehan, M. Imran Nisar, Sana Syed).

**Keywords:** environmental enteropathy, environmental enteric dysfunction, genome-scale metabolic model, pediatric, growth restriction

## Abstract

Environmental Enteric Dysfunction (EED) is a leading contributor to childhood stunting. Most EED literature derives from postnatal pediatric cohorts, leaving unclear how much of the metabolic signature reflects acquired intestinal pathology versus growth restriction originating earlier. We applied genome-scale metabolic modeling to explore whether metabolic features of pediatric EED overlap with those of maternal plasma proteomes associated with adverse early-life growth. Duodenal transcriptomics from the Study of Environmental Enteropathy and Malnutrition (SEEM; 52 children with EED, 24 controls) and maternal plasma proteomics from the Alliance for Maternal and Newborn Health Improvement (AMANHI; n = 2,485 pregnant women) were integrated with the Human1 model using Reaction Inclusion by Parsimony and Transcript Distribution (RIPTiDe), followed by flux sampling and Linear Discriminant Analysis, across five growth-related proxies (small-for-gestational-age, low birth weight, preterm birth, stunting, and wasting). Amino acid and nucleotide metabolism were prominent among the top discriminating reactions in EED and most proxies. Of the 50 reactions distinguishing EED from control, 24 (48%) were among the top reactions in at least one proxy, and 19 (38%) with low birth weight, the largest of any proxy. These exploratory findings suggest shared metabolic pathways across tissues and life stages and motivate longitudinal studies.

## Introduction

Growth restriction in early life remains among the most pressing global health challenges, contributing to substantial child mortality and developmental impairment, with the greatest burden concentrated in South Asia and sub-Saharan Africa (1). Environmental Enteric Dysfunction (EED) is a subclinical disorder of the small intestine, characterized by chronic mucosal inflammation, blunted villous architecture, and increased intestinal permeability, that is highly prevalent among children in low- and middle-income countries with inadequate water and sanitation, and is increasingly recognized as an important contributor to childhood stunting (2).

Much of what is known about EED comes from infant and early-childhood growth studies conducted largely in isolation from antenatal and maternal growth data. Cohorts are typically enrolled postnatally at ages ranging from roughly 12 months to under 5 years, and growth faltering is assessed relative to chronological age with limited integration of pregnancy-level exposures such as small-for-gestational-age (SGA) status, preterm delivery, or intrauterine growth restriction (IUGR). This is a meaningful gap: current clinical practice explicitly corrects anthropometric assessment for gestational age up to 24 months, since growth status calculated from chronological rather than corrected age misclassifies a substantial proportion of preterm infants as stunted or underweight (3). Because most EED cohorts do not systematically account for this antenatal growth trajectory, it remains unclear how much of the postnatal "EED signal" reported in the literature reflects an independent, acquired intestinal pathology versus growth restriction that originated in utero and simply persists into the window where EED is typically studied.

This gap is particularly highlighted in recent multi-country efforts to characterize EED at the molecular level. The Environmental Enteric Dysfunction Biopsy Initiative (EEDBI) Consortium has provided a cross-country appreciation of mucosal and molecular EED signatures spanning Pakistan, Zambia, and Bangladesh, and integrative multi-omics work using machine learning has further characterized immunologic and metabolic dysregulation in pediatric EED across these same populations (4,5). Duodenal transcriptomic profiling from the Study of Environmental Enteropathy and Malnutrition (SEEM) cohort specifically has implicated altered lymphocyte activation and lipid metabolism pathways in Pakistani children with EED, and converging clinical and mechanistic evidence suggests that essential amino acid sufficiency, and tryptophan metabolism in particular, may be a pathway linking poor access to animal-source protein with both EED and stunted growth (6–9). Yet despite this substantial cross-country body of work, these studies were not designed to determine how much of the EED signal they describe may be attributable to maternal-fetal factors- resulting in outcomes such as SGA, preterm birth, or IUGR- that predate and could predispose to EED, as opposed to reflecting EED as an independent postnatal process.

In silico metabolic modeling is a computational method for modeling the relationships between genes, proteins, and chemical reactions within an organism’s cells. A Genome-scale Metabolic Model (GEM) is a generic mathematical model of the cell, permitting a large number of simulations that provide a systems-level view of the organism’s chemical biology. Unlike traditional methods such as Differential Expression Analysis (DEA) or Pathway Enrichment Analysis, metabolic modeling can generate hypotheses in the form of fine-grained reaction-level differences. Our work explores whether metabolic features of maternal plasma associated with adverse early-life growth outcomes overlap with those of biopsy-characterized pediatric EED, as a step toward addressing the maternal-fetal gap in the existing postnatal EED literature. Here, we developed a multi-omics analysis pipeline to identify metabolic features in pregnant mothers whose children experienced adverse growth outcomes and compared them with metabolic features of children with known EED. Because the two cohorts are independent and not participant-linked, this analysis is exploratory.

## Methods

### Study Design and Overview

This study utilized data from two (separately funded) existing cohorts to explore metabolic features associated with EED and early-life growth outcomes (**Figure 1A**). The cohorts are independent and not participant-linked. Cohort A (Alliance for Maternal and Newborn Health Improvement or AMANHI) is a mother-infant cohort from the AMANHI biorepository, a multi-center, population-based cohort study from 2014 to 2018 across five sites in South Asia and six sites in sub-Saharan Africa (10). Cohort B drew on the Study of Environmental Enteropathy and Malnutrition (SEEM), a prospective cohort study of pediatric patients evaluated for EED at the Aga Khan University, Karachi, Pakistan, between 2016 and 2019 (11). Further details are provided below and in the supplementary methods.

**Figure 1.**
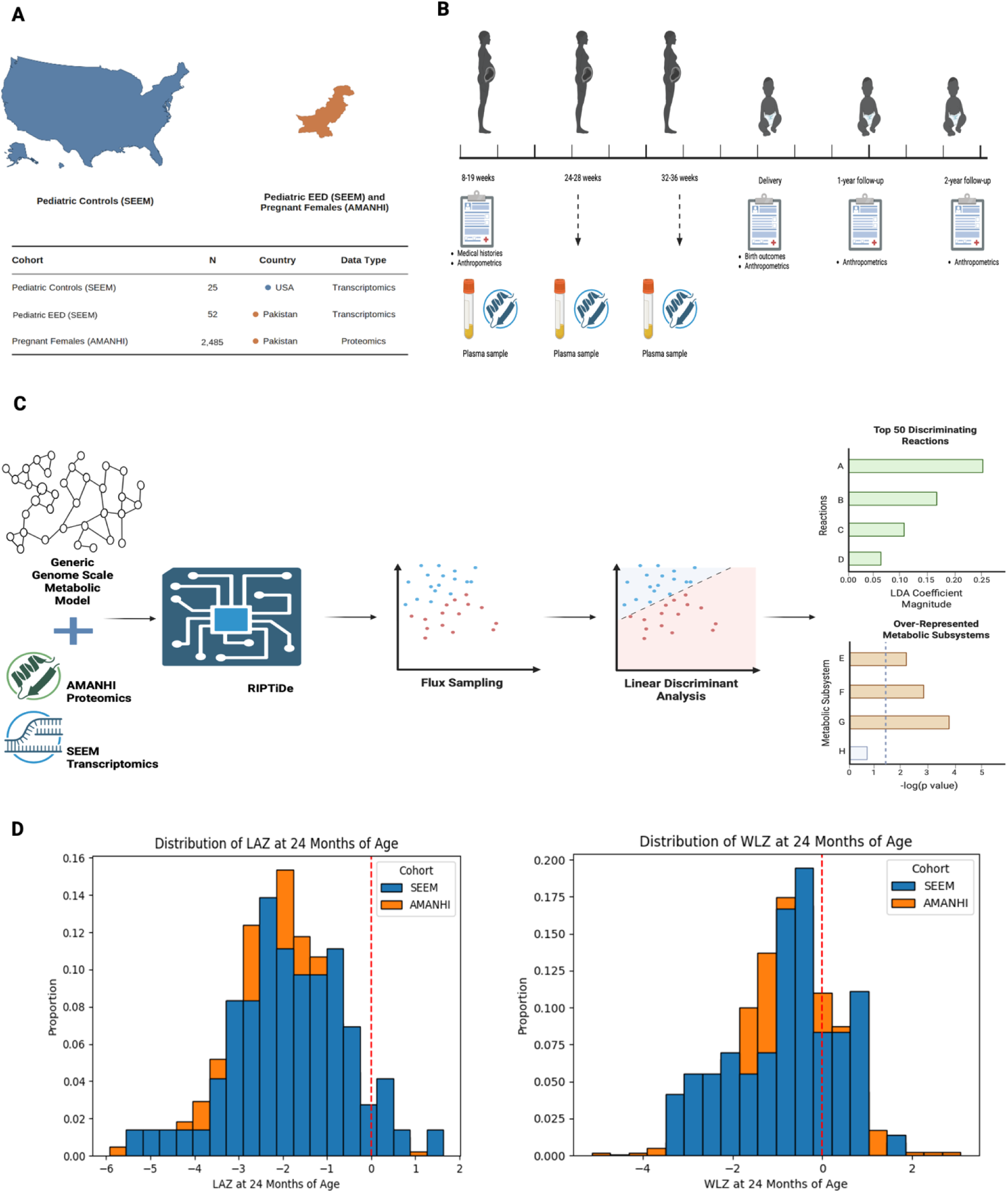
Study design, analytical workflow and growth outcomes. (A) Geographic distribution of study cohorts, including the Study of Environmental Enteropathy and Malnutrition (SEEM) (1) control cohort (n = 24, United States, transcriptomics), the SEEM Environmental Enteric Dysfunction (EED) cohort (n = 52, Pakistan, transcriptomics), and Alliance for Maternal and Newborn Health Improvement (AMANHI) (2) participants (n = 2,485, Pakistan, proteomics) local to the Aga Khan University. (B) Study timeline showing maternal and infant visits across pregnancy (8–19, 24–28, and 32–36 weeks gestation) through delivery and follow-up (1-year and 2-year visits). Plasma samples were collected at each antenatal visit, with medical histories and anthropometrics collected throughout; birth outcomes and anthropometrics were recorded at delivery, and anthropometrics were repeated at the 12- and 24-month follow-ups. (C) Metabolic modeling workflow: a generic genome-scale metabolic model (Human Genome-scale Metabolic Model; Human1) was integrated with participant-level AMANHI proteomics or SEEM transcriptomics data using Reaction Inclusion by Parsimony and Transcript Distribution (RIPTiDe) (3) to generate context-specific metabolic models. Flux sampling was performed on the resulting models, followed by Linear Discriminant Analysis (LDA) and subsequent Fisher Exact Tests to identify the top discriminating reactions distinguishing comparison groups and over-represented metabolic subsystems. P-values were adjusted using the Benjamini-Hochberg procedure with FDR <0.05 as the threshold for significance. (D) Distribution of length-for-age Z-score (LAZ) at 24 months of age among SEEM and AMANHI participants, illustrating the range of linear growth status captured across both cohorts. (E) Distribution of weight-for-length Z-score (WLZ) at 24 months of age among SEEM and AMANHI participants, illustrating the range of weight-for-length status captured across both cohorts. Abbreviations: SEEM = Study of Environmental Enteropathy and Malnutrition; EED = Environmental Enteric Dysfunction; AMANHI = Alliance for Maternal and Newborn Health Improvement; Human1 = Human Genome-scale Metabolic Model; RIPTiDe = Reaction Inclusion by Parsimony and Transcript Distribution; LDA = Linear Discriminant Analysis; FDR = False Discovery Rate; LAZ = length-for-age Z-score; WLZ = weight-for-length Z-score; N = number (sample size); USA = United States of America.

### Study Populations

#### Cohort A (AMANHI)

Cohort A was drawn from the AMANHI biorepository, restricted to participants enrolled from Ibrahim Hyderi and Rerhi Goth, two urban slums in Karachi, Pakistan. Pregnant women were enrolled via field visits conducted by trained Community Health Workers (CHWs); a total of 2,485 women were enrolled (**Figure 1B**). Data were collected at three time points: 8-19 weeks, 24-28 weeks and 32-36 weeks of gestation. Maternal clinical meta-data collected included demographics, socioeconomic status, anthropometrics, medical and obstetric histories, pregnancy, delivery and post-natal birth outcomes. Infant anthropometrics were measured at birth, 1 and 2 years of age.. Maternal and infant anthropometric measurements were collected by trained fieldworkers using standardized, harmonized protocols implemented across all AMANHI sites (12).

#### Cohort B (SEEM)

Cohort B was drawn from the SEEM, a multi-institutional collaboration between the Aga Khan University Hospital (AKUH) and Cincinnati Children’s Hospital Medical Center (CCHMC), conducted at a field site in Matiari, Pakistan. SEEM enrolled a longitudinal birth cohort of 350 children with moderate-to-severe malnutrition (weight-for-length Z-score [WLZ] < −2) and 50 well-nourished children (WLZ > 0 and length-for-age Z-score > −1 on two consecutive visits) between 0 and 6 months of age. Enrolled children received staged educational and nutritional interventions; those who remained malnourished (WLZ < −2) despite intervention underwent medical evaluation to identify a treatable secondary cause of malnutrition, including, where clinically indicated, upper gastrointestinal (UGI) endoscopy with duodenal biopsy.

### Sample Collection

#### Cohort A (AMANHI)

Cohort A collected maternal blood, urine, and stool samples, placental tissue, and infant cord blood (12); only maternal antenatal plasma (for proteomic profiling) and associated clinical and anthropometric metadata were used in this study. See supplemental methods for details of sample collection.

#### Cohort B (SEEM)

Cohort B participants who failed to respond to nutritional rehabilitation and had no identifiable cause of malnutrition on initial laboratory workup underwent upper gastrointestinal (UGI) endoscopy at the Aga Khan University Hospital (11). Endoscopies were performed by a trained pediatric gastroenterologist under sedation, and duodenal biopsies were obtained and utilized for bulk RNA sequencing. The subset with duodenal transcriptomic data available comprised the analysis population for Cohort B: 52 children with EED enrolled in Pakistan and 24 comparator controls with normal duodenal histopathology enrolled in the United States. See supplemental methods for details of sample collection.

### Definition of Growth and Nutritional Status Outcome Groups (EED Proxy Phenotypes)

In Cohort B (SEEM), EED status was established directly, based on duodenal tissue histology obtained via upper gastrointestinal endoscopy. In Cohort A (AMANHI), no equivalent direct measure of EED was available. AMANHI collected extensive antenatal data (absent from the SEEM cohort, which lacked antenatal measures entirely), but did not include tissue-based diagnosis of EED. Because EED was not directly measured in AMANHI, Cohort A participants could not be classified into confirmed EED and non-EED groups as was possible for Cohort B. We therefore examined five measurable, malnutrition-associated phenotypes that served as growth-related proxies rather than direct measures of EED: pre-term birth, small-for-gestational-age, low birth weight, stunting at 2 years of age, and wasting at 2 years of age. These five proxy phenotypes were selected to capture impaired growth and malnutrition-related outcomes; they are not equivalent to EED. See supplemental methods for definitions for each of the five proxy phenotypes.

### Biospecimens

#### Cohort A (AMANHI) Biomarkers

A panel of maternal biomarkers measured from blood samples collected during antenatal visits included general hematologic and metabolic markers- hemoglobin, hemoglobin A1c (HbA1c), calcium, and ferritin; renal and hepatic function markers- blood urea nitrogen (BUN), serum creatinine (sCr), aspartate aminotransferase (AST), and albumin (ALB); infectious and inflammatory markers- C-reactive protein (CRP) and rapid plasma reagin (RPR); and an endocrine marker, thyroid stimulating hormone (TSH).

#### Cohort A (AMANHI) Plasma Proteomics

A total of 100 μL of maternal plasma per sample (two aliquots per participant) was shipped from Aga Khan University, Pakistan, to the Max Planck Institute of Biochemistry, Germany, for untargeted proteomic analysis. From each sample, 5 μL of plasma was processed using a perchloric acid workflow with neutralization (PCA-N), which significantly reduces costs with greater proteomic depth (13). See supplemental methods for full proteomics workflow.

#### Cohort B (SEEM) Duodenal Tissue Transcriptomics

RNA was extracted from duodenal biopsy tissue collected at endoscopy from 52 children with Environmental Enteric Dysfunction (EED) and 24 comparator controls. Sequencing libraries were prepared using polyA RNA selection, followed by fragmentation, complementary DNA (cDNA) synthesis, and adaptor ligation, using the TruSeq RNA Sample Preparation kit (Illumina) (6,11). Raw sequencing reads (FASTQ files) were processed using a standardized, cloud-based RNA-sequencing pipeline and gene-level transcript counts were subsequently quantified for use in downstream context-specific metabolic modeling. See supplemental methods for full transcriptomics workflow.

### Genome-Scale Metabolic Modeling

#### Model Integration and Flux Sampling

Context-specific metabolic models were constructed for each comparison subcohort by integrating a generic genome-scale metabolic model (Human1) with cohort-averaged ’omics data- AMANHI maternal plasma proteomics or SEEM duodenal tissue transcriptomics, as appropriate to the cohort (**Figure 1C**). Quantified proteins or transcripts were mapped to their corresponding genes and reactions within Human1. Reaction Inclusion by Parsimony and Transcript Distribution (RIPTiDe) was then applied to generate context-specific models: omics-derived abundance measures were used to weight reaction likelihoods, such that reactions associated with higher molecular abundance were preferentially retained during model pruning, while overall network flux was minimized to yield a metabolically feasible and parsimonious network without reliance on arbitrary abundance cutoffs. All default parameters for RIPTiDe were preserved, except the minimum objective fraction (increased from 0.8 to 1.0). To characterize the range of feasible metabolic states for each context-specific model, flux sampling was performed to uniformly and efficiently interrogate the constrained solution space. These flux samples represented plausible metabolic states within each subcohort and were used as the basis for identifying reaction-level differences between comparison groups.

Because models were built from cohort-averaged data, sampled flux states describe each model’s solution space, not independent participant-level observations.

#### Identification of Discriminating Reactions and Over-Represented Subsystems

Linear Discriminant Analysis (LDA) was applied to the flux samples from each pair of context-specific models to identify the axis of maximal separation between comparison groups. To meet LDA assumptions, sampled flux was transformed via quantile transformation and multicollinearity was addressed using the Ledoit-Wolf shrinkage estimator. Reactions were ranked by the magnitude of their LDA coefficient, and the top 50 reactions with the largest coefficients were retained as the discriminating reactions for each comparison. For each comparison, the 10 reactions with the highest-magnitude LDA coefficients (of the top 50 identified) were selected for detailed flux-level characterization. Sampled flux distributions for these 10 reactions were compared between comparison groups using a two-sided

Mann-Whitney U test, with a significance threshold of 0.05. These tests describe differences between model-derived flux distributions, not population-level inference. The metabolic subsystem membership of each of these top 10 reactions was independently determined by expert human clinical and biochemical review based on the reaction’s stoichiometric equation, substrate/product identity, and known metabolic pathway context.

#### Cross-Cohort Comparison of Discriminating Reactions and Subsystems

To assess whether metabolic signatures identified in pediatric EED were reflected in early-life proxy phenotypes, the top 50 discriminating reactions from the SEEM EED vs. Control comparisons were compared against the top 50 discriminating reactions from each of the five AMANHI proxy comparisons: SGA vs. AGA, LBW vs. NBW, pre-term vs. term, stunting vs. no stunting, and wasting vs. no wasting. Reactions were considered shared between comparisons if they were identified as top discriminating reactions in both the SEEM EED comparison and one or more AMANHI proxy comparisons. Shared reactions and their associated subsystems were evaluated both at the level of individual reactions and at the level of broader metabolic pathway groupings across cohorts.

### Statistical and Bioinformatic Analysis

Two-group comparisons were conducted using the Mann-Whitney U-test or the Kolmogorov-Smirnov test (14,15). The former was applied when testing for a difference in medians between distributions. The latter was applied to determine whether two distributions significantly differed in shape (even if their measures of central tendency were similar).

Association between binary categorical variables was determined by Fisher’s Exact Test, as the tested categories were often rare (making the χ² Test asymptotic approximations unreliable) (16). When the number of categories in one or both variables exceeded 2, a Monte-Carlo approximation of Fisher’s Exact Test was employed with 10 million rollouts. Multiple hypotheses were corrected using the Benjamini-Hochberg method when prioritizing hypothesis generation and preserving statistical power, or the Bonferroni method when prioritizing minimal false-positive results (17,18). See supplemental methods for complete statistical and bioinformatics analysis.

### Ethical Approval and Consent

See supplemental methods

### Data and Code Availability

Transcriptomic data are available at GEO (GSE159495) and analysis code at https://github.com/SyedLab-GI/CompEE; AMANHI proteomic data may be requested from FJ and IN. See supplemental methods.

## Results

### Cohort Characteristics

#### Cohort A (AMANHI) Demographics and Clinical Characteristics

A total of 2,485 pregnant women were enrolled in Cohort A from the Karachi, Pakistan sites of the AMANHI biobank. The median maternal age at enrollment was 26 years (IQR: 23–30), with a median height of 153.7 cm (IQR: 150.0–158.0) and median pre-pregnancy BMI of 21.6 kg/m² (IQR: 18.8–25.3) (**Table 1 A)**. By BMI category, 52.4% of women were classified as healthy weight (18.5–25 kg/m²), 21.1% underweight (< 18.5 kg/m²), 18.9% overweight (25–30 kg/m²), 7.2% obese (30–40 kg/m²), and 0.4% morbidly obese (> 40 kg/m²). Maternal biomarker screening identified anemia (hemoglobin < 11.6 g/dL) in 62.7% of women, elevated CRP (≥ 0.9 mg/dL) in 26.4%, and abnormal HbA1c (> 5.6%) in 3.3%.

**Table 1.**
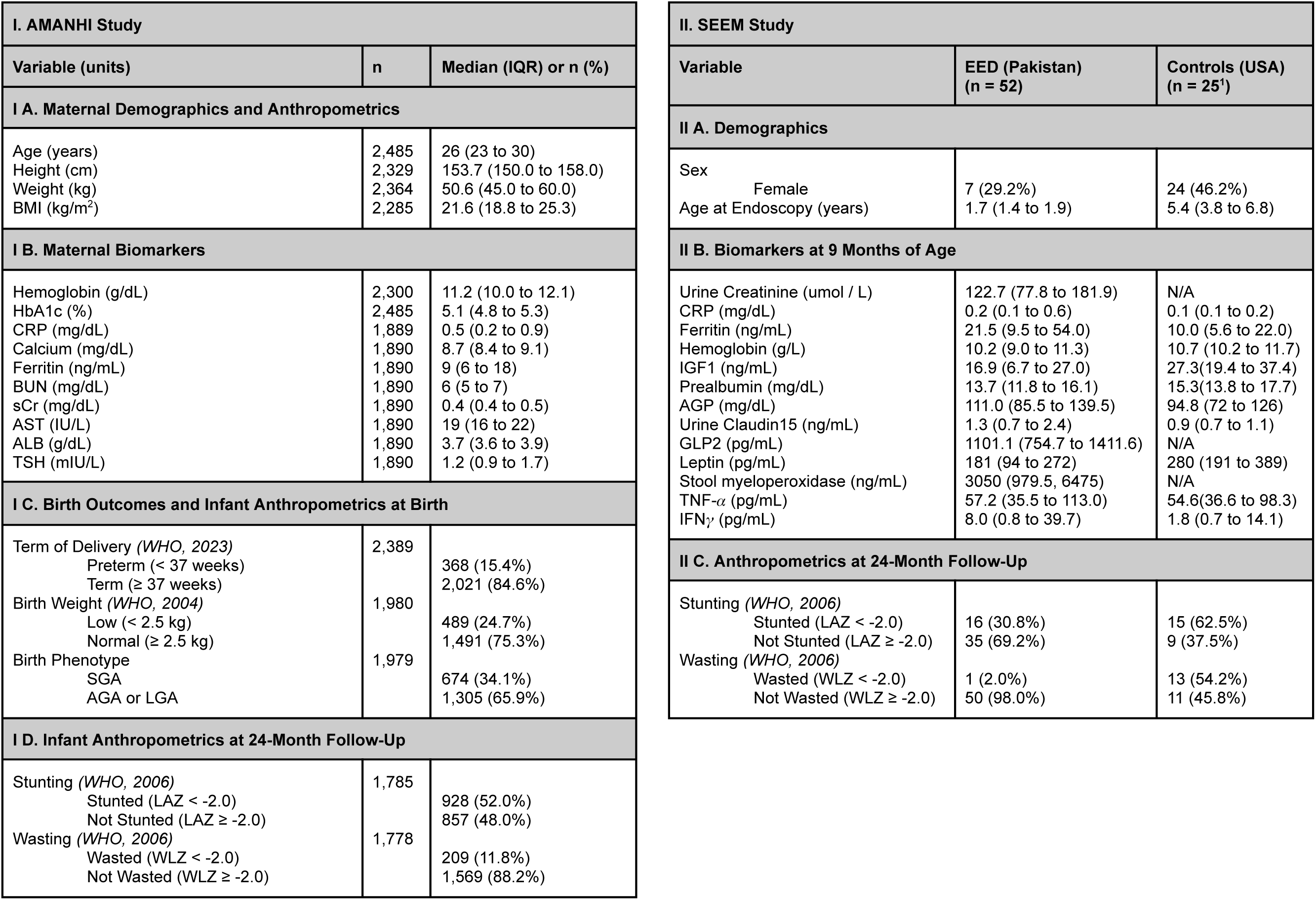
Descriptive summary of (I) AMANHI and (II) SEEM cohorts. In table IA, variables are grouped by (A) maternal demographics and anthropometrics, (B) maternal biomarkers, (C) birth outcomes and infant anthropometrics at birth, and infant anthropometrics at (D) 24-month follow-ups. For each variable, the median and inter-quartile range are provided if the variable is numerical, the number and percentage of participants in the minority are indicated if the variable is binary, and the number and percentage of participants for each category if the variable is categorical. Not all variables were available for all participants; the number of participants each variable was measured for is recorded in the middle column. In table B, variables are grouped by (A) demographics, (B) biomarkers at 9 months of age, and (C) anthropometrics at 24-month follow-up. For each variable, the median and inter-quartile range are provided if the variable is numerical, the number and percentage of participants in the minority are indicated if the variable is binary, and the number and percentage of participants for each category if the variable is categorical. Note that not all variables were available for all participants and that many variables were not collected for the CCHMC control subcohort of the SEEM study. ^1^1 of 25 control patients did not consent to their transcriptomic data being publicly shared. Although this patient was included for Table 1, they were excluded for all future analysis. AMANHI = Alliance for Maternal and Newborn Health Improvement; SEEM = Study of Environmental Enteropathy and Malnutrition; CCHMC = Cincinnati Children’s Hospital Medical Center; BMI = body mass index; HbA1c = hemoglobin A1c; CRP = c-reactive protein; BUN = blood urea nitrogen; sCr = serum creatinine; AST = aspartate aminotransferase; ALB = albumin; TSH = thyroid stimulating hormone; WHO = World Health Organization; SGA = small for gestational age; AGA = appropriate for gestational age; LGA = large for gestational age; LAZ = length-for-age Z-score; WAZ = weight-for-age Z-score; WLZ = weight-for-length Z-score; IGF1 = insulin-like growth factor 1; AGP = alpha-1-acid glycoprotein; GLP2 = glucagon-like peptide-2; TNF-*α* = tumor necrosis factor alpha; IFN*γ* = interferon-gamma

Among 2,461 pregnancies with a recorded outcome, 2,323 (94.4%) resulted in a live birth, 91 (3.7%) in stillbirth, and 47 (1.9%) in miscarriage. Of 2,389 deliveries with gestational age recorded, 84.6% occurred at term (≥ 37 weeks), while 15.4% were pre-term. Infants were 48.5% female (n = 2,318 with sex recorded). At birth, 24.8% of infants were classified as low birth weight (< 2.5 kg), with a median birth weight percentile of 17.7% (IQR: 6.3–37.7). Based on birth weight percentile, 34.1% of infants (675/1,979) were classified as small-for-gestational-age (SGA) and 65.9% (1,304/1,979) as appropriate-for or large-for-gestational-age (AGA or LGA).

Median length-for-age (LAZ), weight-for-age (WAZ), and weight-for-length (WLZ) Z-scores at birth were −0.8 (IQR: −1.6 to 0.0), −1.2 (IQR: −1.9 to −0.6), and −1.0 (IQR: −1.9 to −0.2), respectively (**Figure 1D**).

#### Cohort B (SEEM) Demographics and Clinical Characteristics of Children with EED and Comparator Controls

Cohort B comprised 52 children with Environmental Enteric Dysfunction (EED) enrolled in Pakistan and 24 comparator controls enrolled in the United States (**Table 1 B)**. Children with EED underwent endoscopy at a markedly younger median age (1.7 years, IQR: 1.4 -- 1.9) than controls (5.4 years, IQR: 3.8 -- 6.8), reflecting the clinical indication for endoscopy in the malnourished SEEM cohort. At the time of endoscopy, children with EED showed substantially lower anthropometric z-scores than controls: median LAZ of −3.2 (IQR: −3.6 to 2.3) versus 0.1 (IQR: −0.5 to 0.8) in controls, and median WAZ of −2.9 (IQR: −3.5 to −2.6) versus −0.1 (IQR: −1.1 to 0.8) in controls.

### Context-Specific Metabolic Modeling Highlights Amino Acid and Nucleotide Metabolism Among Discriminating Subsystems in EED and SGA

Context-specific genome-scale metabolic models were successfully constructed for all six comparisons: SEEM EED vs. Control (duodenal transcriptomics) and the five AMANHI proxy comparisons- SGA vs. AGA, LBW vs. NBW, preterm vs. term, stunting vs. no stunting, and wasting vs. no wasting (maternal plasma proteomics). For each comparison, omics-derived gene/protein abundance data were integrated with the generic Human1 reconstruction using RIPTiDe, yielding a pair of context-specific models per phenotype. Approximately 500-1,000 reactions were retained after pruning, varying by context-specific model. One thousand feasible metabolic states were sampled for each model, yielding a distribution of flux for each reaction in the context-specific model. When comparing two context-specific models (e.g. SGA vs AGA), only reactions shared between them were considered. We applied Linear Discriminant Analysis (LDA) to identify the reactions whose sampled flux best separated each comparison group. For each of the six comparisons, the 50 reactions with the highest-magnitude LDA coefficients were obtained. To determine the enriched pathways, we A) grouped the top 50 reactions using the subsystems intrinsic to HumanGEM and B) grouped the top 10 reactions by expert clinical review. The former method was followed by a Fisher’s Exact Test with Benjamini-Hochberg corrected p-values to identify significantly (FDR < 0.05) enriched pathways. The latter method was applied primarily for illustrative purposes – quantitative conclusions were not sought.

The top 10 reactions differentiating EED from control belonged to the Pentose Phosphate Pathway (ATP:D-ribose 5-phosphotransferase, exchange of ribose, MAR04958, D-Ribose 1,5-phosphomutase), Nucleotide Metabolism (inosine:phosphate alpha-D-ribosyltransferase, MAR05042, exchange of inosine), Redox Metabolism (NADPH:NAD+ oxidoreductase), Amino Acid Metabolism (exchange of glutamate), and Fatty Acid Synthesis (Stearoyl Coenzyme A Desaturase) (**Figure 2A**). The same approach applied to the AMANHI SGA vs. AGA comparison identified the top reactions as belonging to Cholesterol Metabolism (exchange of LDL, exchange of LDL remnant, MAR00014), Nucleotide Metabolism (ATP:dTDP phosphotransferase), Vitamin A Metabolism (retinal:NAD+ oxidoreductase, MAR06650), Amino Acid Metabolism (MAR05330, MAR09605), and Fatty Acid Oxidation (Hexanoyl-CoA hydrolase, (S)-Hydroxyoctanoyl-CoA hydro-lyase) (**Figure 2B**).

**Figure 2.**
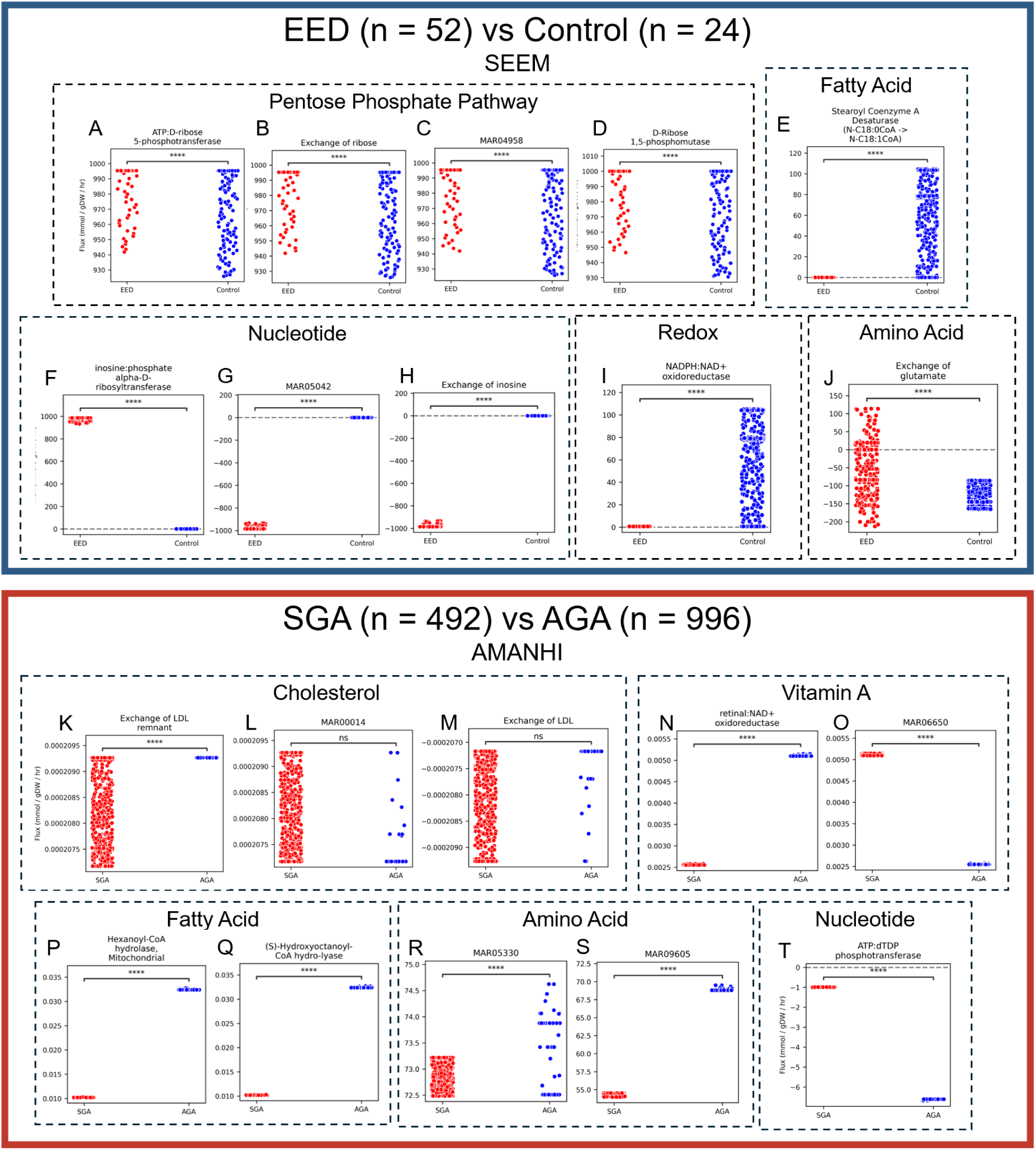
Altered Nucleotide Metabolism, Amino Acid Metabolism, and Fatty Acid-Related Pathways Differentiate EED from Controls and SGA from AGA. Jitter plots showing the top 10 reactions identified by our Reaction Inclusion by Parsimony and Transcript Distribution (RIPTiDe) and Linear Discriminant Analysis framework. Our framework was applied to SEEM duodenal transcriptomics (A-J) and to AMANHI maternal plasma proteomics from visit 2 (24-28 weeks) (K-T). The top 10 reactions were categorized into subsystems based on each reaction’s biochemical function by literature review. In each plot, the x-axis describes the altered reaction between 1) EED (n = 52) or 2) SGA (n = 492) and 1) controls (n = 24) or 2) AGA (n = 996). The y-axis shows the flux values sampled from the context-specific metabolic network reconstructions produced by RIPTiDe. A two-sided Mann-Whitney U-test was used to assess the significance of differences in flux distributions. Note that the scale of the y-axes between plots varies and is shown primarily for illustrative purposes. Both sets of discriminating reactions implicate Nucleotide Metabolism (F, G, H, T), Amino Acid Metabolism (J, R, S), and Fatty Acid-related pathways (E, P, Q). n.s. = not significant (p >= 0.05) * = 0.01 < p < 0.05 ** = 0.001 < p < 0.01 *** = 0.0001 < p < 0.001 **** = 0.00001 < p < 0.0001 Abbreviations: EED = Environmental Enteric Dysfunction; SEEM = Study of Environmental Enteropathy and Malnutrition; SGA = Small for Gestational Age; AGA = Appropriate for Gestational Age; AMANHI = Alliance for Maternal and Newborn Health Improvement; MAR04958 = ribose [c] ⇔ ribose [e]; MAR05042 = inosine [c] ⇔ inosine [e]; MAR00014 = LDL [e] ⇒ LDL remnant [e] + 680 cholesterol [e] + 1515 cholesterol-ester pool [e]; MAR06650 = retinal [r] ⇒ 9-cis-retinal [r]; MAR05330 = Na^+^ [e] + glutamate [e] ⇒ Na^+^ [c] + glutamate [c]; MAR09605 = H^+^ [c] + Na^+^ [e] + alanine [e] ⇒ H^+^ [e] + Na^+^ [c] + alanine [c]; ATP = adenosine triphosphate; NADPH = nicotinamide adenine dinucleotide phosphate; NAD+ = nicotinamide adenine dinucleotide; LDL = low density lipoprotein; dTDP = thymidine diphosphate; Na^+^ = sodium ion; H^+^ = hydrogen ion

Despite being derived from independent cohorts, tissue types, and omics modalities, the top discriminating reactions for EED vs. Control and SGA vs. AGA converged on three shared metabolic subsystems: Nucleotide Metabolism, Amino Acid Metabolism, and Fatty Acid-related pathways, suggesting these pathways as shared metabolic features distinguishing both phenotypes from their respective comparison groups. Similar comparison groups with EED were constructed for stunting vs no stunting and LBW vs NBW **(Supplemental figures 1 and 2).**

### Cross-Cohort Comparison Identifies Overlapping Metabolic Reactions Between EED and Early-Life Growth Phenotypes

Having identified the top reactions distinguishing EED from control and each AMANHI proxy phenotype from its comparison group individually, we next asked whether reactions distinguishing pediatric EED also distinguished the early-life growth-related proxies derived from maternal plasma. We compared the top 50 reactions distinguishing SEEM EED from Control against the top 50 reactions distinguishing each of the five AMANHI proxy comparisons (**Figure 3**). Of the 50 reactions identified in the EED vs. Control comparison, 24 (48%) were also identified among the top 50 discriminating reactions in at least one AMANHI proxy comparison. D-glyceraldehyde-3-phosphate aldose-ketose-isomerase (Central Carbon Metabolism) was the most widely shared reaction, identified in EED and in all five AMANHI proxy comparisons.

**Figure 3.**
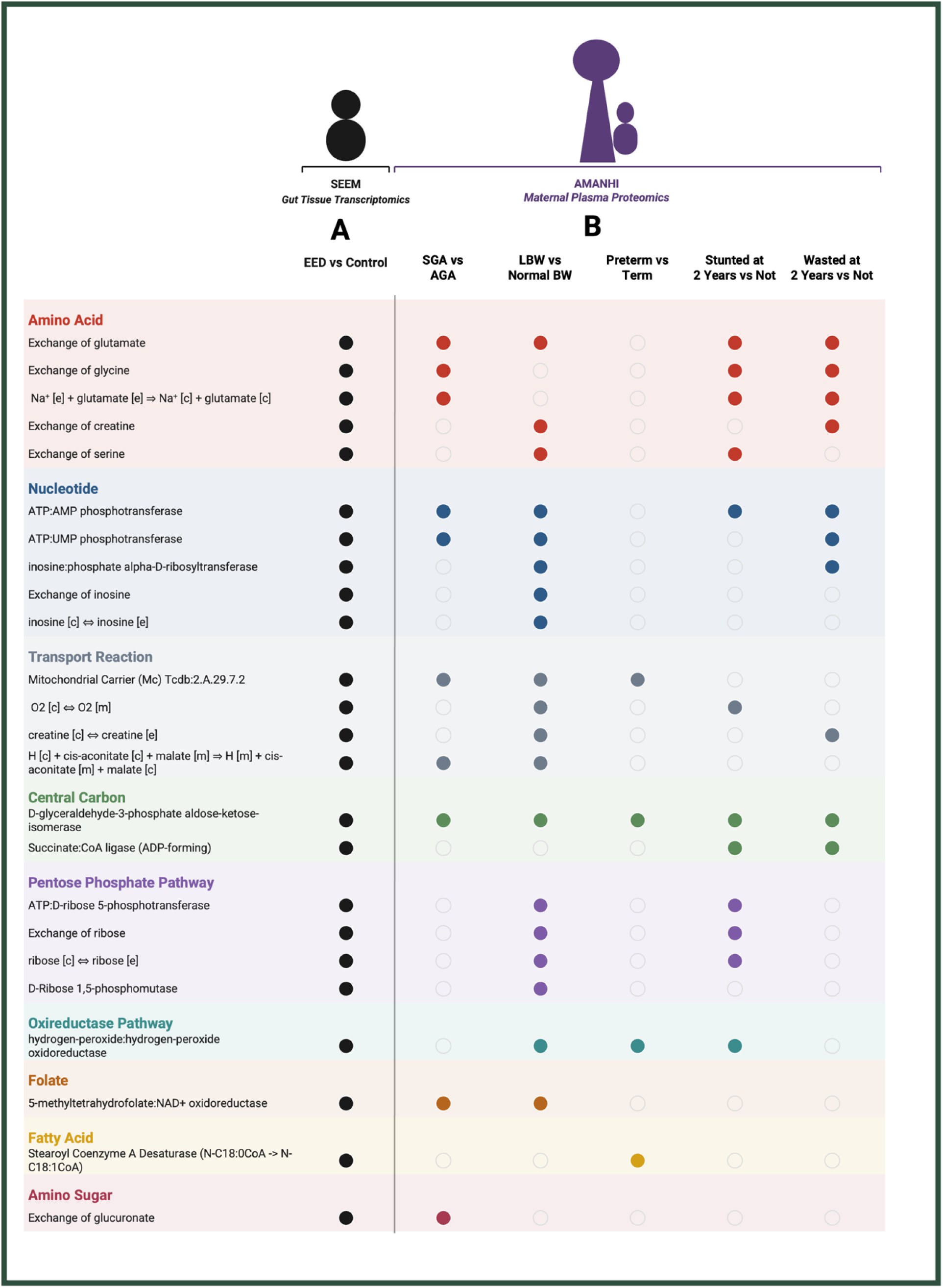
Amino Acid and Nucleotide Metabolism Shows Shared Signals Between Pediatric EED and Maternal-Fetal Growth Restriction Proxy Cohorts. Dot-matrix plot showing the 24 reactions (of the top 50 reactions distinguishing the Study of Environmental Enteropathy and Malnutrition (SEEM) (1) EED cohort from controls) that were also among the top discriminating reactions in at least one of five EED proxy comparisons drawn from the Alliance for Maternal and Newborn Health Improvement (AMANHI) (2) cohort: small-for-gestational-age (SGA) vs. appropriate-for-gestational-age (AGA), low birth weight (LBW) vs. normal birth weight (NBW), pre-term vs. term delivery, stunting vs. no stunting at 2 years, and wasting vs. no wasting at 2 years. Column ‘A’ (black-filled circles) represents the SEEM EED reference cohort (pediatric duodenal tissue), from which all 24 reactions were originally identified; ‘B’ represent the AMANHI proxy comparisons (maternal plasma proteomics collected during pregnancy, with offspring later classified into each proxy group at birth or at 2-year follow-up). A filled circle indicates the reaction was among that group’s top 50 discriminating reactions; an open circle indicates it was not. Reactions are grouped into nine metabolic subsystems (color-coded, left margin): Amino Acid Metabolism, Nucleotide Metabolism, Transport Reaction, Central Carbon Metabolism, Pentose Phosphate Pathway, Oxireductase Pathway, Folate Metabolism, Fatty Acid Metabolism, and Amino Sugar Metabolism. SEEM: Study of Environmental Enteropathy and Malnutrition; EED: Environmental Enteric Dysfunction; AMANHI: Alliance for Maternal and Newborn Health Improvement; SGA: small-for-gestational-age; AGA: appropriate-for-gestational-age; LBW: low birth weight; NBW: normal birth weight.

Exchange of glutamate (Amino Acid Metabolism) and ATP:AMP phosphotransferase (Nucleotide Metabolism) were each shared between EED and four of the five proxy comparisons. Examining all 5 of the AMANHI proxies, we noted that LBW vs. NBW shared the largest number of most discriminating reactions with EED (19 of the 50 EED reactions, 38%; 79% of the 24 shared reactions), followed by stunting (12 of 50, 24%; 50% of shared reactions), while SGA vs. AGA and wasting each shared 10 of 50 (20%; 42% of shared reactions), and pre-term birth shared only 4 of 50 (8%; 17% of shared reactions).

These 24 shared reactions spanned nine metabolic subsystems: Amino Acid Metabolism, Nucleotide Metabolism, Transport Reactions, Central Carbon Metabolism, Pentose Phosphate Pathway, the Oxireductase Pathway, Folate Metabolism, Fatty Acid Metabolism, and Amino Sugar Metabolism. Amino Acid Metabolism and Nucleotide Metabolism accounted for the largest number of shared reactions between EED and the AMANHI proxies (five each), while Central Carbon Metabolism, though represented by only two reactions, contained the single reaction conserved across all five proxy comparisons.

## Discussion

This study integrated genome-scale metabolic modeling across two independent cohorts to explore whether metabolic features of pediatric EED overlap with those of maternal proteomes associated with adverse early-life growth outcomes. First, reactions from amino acid metabolism, and to a lesser degree nucleotide metabolism, were prominent among the top discriminating reactions in both EED vs Control and across most of the growth-related proxies. Secondly, of the 50 reactions distinguishing EED from Control, nearly half were also identified among the top discriminating reactions in at least one AMANHI proxy comparison. Finally, the greatest overlap with the reactions distinguishing EED was found in the low-birth-weight group from the AMANHI cohort (19 of 50 reactions).

The prominence of amino acid metabolism among these reactions is broadly consistent with prior work linking circulating amino acid depletion to impaired childhood growth (19). In a cohort of Malawian children, stunted children had significantly lower serum concentrations of all nine essential amino acids, along with several conditionally essential and non-essential amino acids including glutamate, whose exchange reaction also emerged among our top discriminating reactions in EED. Semba and colleagues further proposed that amino acid sufficiency governs childhood linear growth through the mechanistic target of rapamycin complex 1 (mTORC1) pathway, which senses intracellular amino acid availability and represses protein synthesis and cellular growth when specific amino acids, particularly glutamine, arginine, and leucine, are deficient (20). Because mTORC1 activity is required for growth in the intestinal epithelium as well as bone and skeletal muscle, amino acid depletion has been proposed as one mechanism that could link impaired systemic growth to intestinal dysfunction, although this was not tested in our analysis.

Relevant to our findings, a separate metabolomic analysis of children with EED identified broad amino acid and lipid metabolic derangements in children with confirmed EED (21). Furthermore, single-cell profiling of duodenal biopsies from children with EED has demonstrated decreased epithelial cell proliferation and reduced goblet cell abundance, both of which were reversible with tryptophan supplementation in vitro (22). Tryptophan is diverted from protein synthesis toward kynurenine by indoleamine-2,3-dioxygenase 1 (IDO1) during chronic inflammation, and the resulting kynurenine-tryptophan ratio has been shown to inversely track with length-for-age Z-score in EED-endemic populations (23). Notably, related findings have been reported in the SEEM cohort itself: mucosal transcriptomic profiling of SEEM duodenal biopsies implicated lymphocyte activation and lipid metabolism pathways in refractory EED, alongside evidence that amino acid supplementation ameliorated villus blunting (6). Taken together, our reaction-level findings and previous literature are compatible with amino acid availability as one candidate link between intestinal dysfunction and systemic growth failure, which our modeling does not establish.

The broader pattern of reaction sharing across EED and all five AMANHI proxy phenotypes is consistent with amino acid and nucleotide metabolism as a convergent feature across these phenotypes. Nearly half of the top 50 EED reactions were also identified among the top discriminating reactions in at least one AMANHI proxy comparison, with D-glyceraldehyde-3-phosphate aldose-ketose-isomerase, a core glycolytic enzyme, as the single most widely shared reaction, appearing among the top discriminators in EED and in all five AMANHI proxies. This overlap was not evenly distributed, however, with low birth weight (LBW) sharing the largest proportion of reactions with EED of any proxy phenotype, followed by stunting, SGA, and wasting. Because LBW also includes preterm infants and preterm birth showed the least overlap, this may indicate a closer relationship with growth restriction than with gestational timing, though these data cannot establish this.

This pattern is consistent with literature linking maternal protein and amino acid status to offspring birth weight. Women consuming higher quantities of animal-source protein during pregnancy deliver infants with higher birth weight and lower risk of fetal growth restriction, and periconceptional animal-source protein intake, unlike plant-source protein, is specifically associated with greater embryonic growth and birth weight in prospective cohort data (24,25,26). Relatedly, a meta-analysis of balanced protein-energy supplementation trials during pregnancy found significantly reduced risk of delivering a growth-restricted infant, with the supplemented group also gaining significantly more gestational weight than controls (27). Given that amino acid metabolism, and glutamate exchange specifically, was among the reactions most broadly shared between EED and the AMANHI proxies, and that LBW showed the largest overlap, a hypothesis for future study is that maternal amino acid and protein adequacy may contribute to both fetal growth restriction and later susceptibility to EED in the resulting infant.

This study has several strengths. First, by pairing biopsy-confirmed pediatric EED (SEEM) with an independent, larger maternal-infant cohort (AMANHI, n = 2,485), we were able to test whether metabolic features identified directly from intestinal tissue also overlap with those from an entirely different biospecimen (maternal plasma), a different omics modality (proteomics versus transcriptomics), and a different population. Second, our use of a shared, standardized genome-scale metabolic modeling pipeline across all comparisons allowed reaction-level and subsystem-level comparison across SEEM and AMANHI. Third, by evaluating five distinct AMANHI growth-restriction proxies rather than a single composite outcome, we observed that the overlap was not uniform across phenotypes and was greatest for LBW, a pattern that would not have been visible with a single-outcome design. Finally, because AMANHI proteomic sampling occurred during pregnancy, this design examines maternal metabolic features measured before the child’s postnatal period, helping to address a gap in a literature that has, to date, characterized EED almost exclusively from postnatal, cross-sectional pediatric sampling.

Our study has several limitations. The AMANHI cohort relies on anthropometric proxies rather than direct EED diagnosis, and these proxies are imperfect indicators of growth restriction: low birth weight captures any small infant, including appropriately grown preterm infants, and therefore does not necessarily reflect impaired fetal growth, in contrast to SGA. The SEEM and AMANHI cohorts are independent, not linked at the participant level, and differ in tissue type, omics modality, geography, and life stage, precluding causal or temporal inference; our findings do not show that maternal metabolic features precede or predict EED. Within SEEM, EED cases (Pakistan) and controls (United States) also differed in age, geography, and nutritional status.

Because models were built from cohort-averaged data, flux comparisons reflect differences between models rather than between individuals. Plasma protein abundance may not reflect intracellular enzyme abundance in any single tissue, primary comparisons were not adjusted for maternal clinical covariates, and the observed overlap was not benchmarked against an empirical null, so highly connected hub reactions may be detected across many unrelated comparisons. Finally, RIPTiDe, originally designed for transcriptomic data, was applied to AMANHI proteomics data (requiring a pre-processing step mapping proteins back to known RNA transcripts), and the altered metabolic pathways lack in vitro validation. These findings should be considered exploratory and hypothesis-generating.

Despite these limitations, the overlap in amino acid and nucleotide metabolism across independently derived cohorts, tissue types, and omics platforms may point to shared metabolic pathways associated with early-life growth restriction and intestinal dysfunction, and warrants further investigation as a candidate for future biomarker or mechanistic studies. Future studies could use cord blood samples from the AMANHI biobank that were not analyzed here; proteomic and metabolomic profiling of cord blood would allow comparison of the fetal metabolic environment at birth against both the maternal antenatal and pediatric EED signatures, which could help clarify whether the pathways highlighted here originate in utero.

## Additional Contributions

We acknowledge the field workers, data management unit and laboratory staff at the Aga Khan University, Karachi, Pakistan, for contributing to the data collection in this work. Anum Chotani, BA and Fatima Zulqarnain, MBBS, from the University of Virginia, Charlottesville, assisted in data coordination for this work. We thank our partners at the Max Planck Institute of Biochemistry for their protoeomic analysis of the plasma samples. We would also like to thank Jason Papin, PhD from the University of Virginia for his support.

## Conflicts of Interest

The authors have no conflicts of interest relevant to this article to disclose.

## Source of Funding

Research reported in this publication was supported by Gates Foundation INV-047735 (SS, IN) and INV-005274 (WK, JK, MFQ, MM, FJ, IN); the National Institutes of Allergy and Infectious Disease (NIAID) K24AI192996 (SS) and National Institute of Diabetes and Digestive and Kidney Diseases (NIDDK) R01DK132369 (SS).

## Ethics

All data used in this study was previously collected in the AMANHI and SEEM studies, obtained with permission. The AMANHI study was approved by the local and institutional ethics committees at all three AMANHI sites, including the Aga Khan University Ethical Review Committee (ERC: 2790-Paeds-ERC-13) for the Pakistan site, and by the World Health Organization Ethics Review Committee (RPC 532). The SEEM study was approved by the Aga Khan University Ethical Review Committee (ERC: 2446 Ped ERC-13) and the Institutional Review Board at Cincinnati Children’s Hospital Medical Center (CCHMC, Study ID 2016-0387), and is registered with ClinicalTrials.gov (NCT03588013).

## Abbreviations

AGA: Appropriate-for-Gestational-Age;
AKUH: Aga Khan University Hospital;
ALB: Albumin;
AMANHI: Alliance for Maternal and Newborn Health Improvement;
AST: Aspartate Aminotransferase;
BMI: Body Mass Index;
BUN: Blood Urea Nitrogen;
CCHMC: Cincinnati Children’s Hospital Medical Center;
cDNA: Complementary DNA;
CHW: Community Health Worker;
CRP: C-Reactive Protein;
DEA: Differential Expression Analysis;
EED: Environmental Enteric Dysfunction;
EEDBI: Environmental Enteric Dysfunction Biopsy Initiative;
ERC: Ethical Review Committee;
FDR: False Discovery Rate;
GEM: Genome-scale Metabolic Model;
HbA1c: Hemoglobin A1c;
Human1: Human Genome-scale Metabolic Model;
IDO1: Indoleamine-2,3-Dioxygenase 1;
IQR: Interquartile Range;
IUGR: Intrauterine Growth Restriction;
LAZ: Length-for-Age Z-score;
LBW: Low Birth Weight;
LDA: Linear Discriminant Analysis;
LGA: Large-for-Gestational-Age;
mTORC1: Mechanistic Target of Rapamycin Complex 1;
NBW: Normal Birth Weight;
PCA-N: Perchloric Acid workflow with Neutralization;
RIPTiDe: Reaction Inclusion by Parsimony and Transcript Distribution;
RPR: Rapid Plasma Reagin;
sCr: Serum Creatinine;
SEEM: Study of Environmental Enteropathy and Malnutrition;
SGA: Small-for-Gestational-Age;
TSH: Thyroid Stimulating Hormone;
UGI: Upper Gastrointestinal;
WAZ: Weight-for-Age Z-score;
WHO: World Health Organization;
WLZ: Weight-for-Length Z-score.

## Contributor’s Statement

The author’s contributions to the paper are as follows: Study concept and design: Sana Syed, Imran Nisar, Fyezah Jehan, Adam Greene Data collection: Waqasuddin Khan, Nathaniel Ribeiro, Shahzaib Khan, Najeeha Talat Iqbal, Asad Ali, Javairia Khalid, M. Farrukh Qazi, M. Mohsin Mannan Analysis and interpretation of results: Nathaniel Ribeiro, Shahzaib Khan, Nazanin Moradinasab, Connor Moore, Lauren J. Klein Draft manuscript preparation: Shahzaib Khan, Nathaniel Ribeiro, Sana Syed All authors have reviewed the results and approved the final version of the manuscript.

---

**Supplementary Figure 1.**
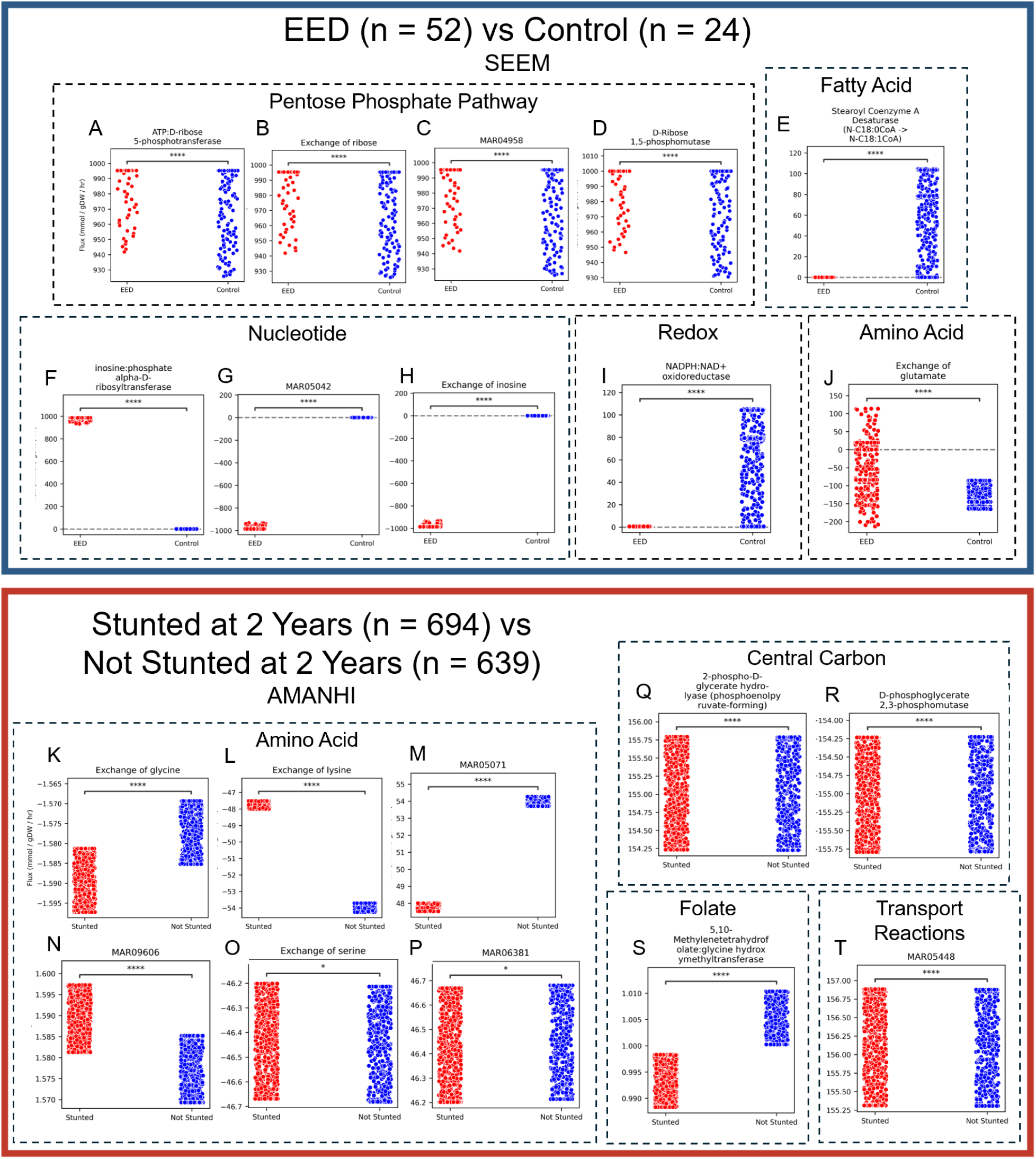
Altered Amino Acid Metabolism Differentiates EED from Controls and Stunted from Not Stunted. Jitter plots showing the top 10 reactions identified by our RIPTiDe and LDA framework. Our framework was applied to SEEM duodenal transcriptomics (A-J) and to AMANHI maternal plasma proteomics from visit 2 (24-28 weeks) (K-T). The top 10 reactions were categorized into subsystems based on each reaction’s biochemical function by literature review.. In each plot, the x-axis describes the altered reaction between 1) EED (n = 52) or 2) Stunted (n = 694) and 1) controls (n = 24) or 2) Not Stunted (n = 639). The y-axis shows the flux values sampled from the context-specific metabolic network reconstructions produced by RIPTiDe. A two-sided Mann-Whitney U-test was used to assess the significance of differences in flux distributions. Note that the scale of the y-axes varies between plots and is shown primarily for illustrative purposes. Both sets of discriminating reactions implicate Amino Acid Metabolism (J, K, L, M, N, O, P). n.s. = not significant (p >= 0.05) * = 0.01 < p < 0.05 ** = 0.001 < p < 0.01 *** = 0.0001 < p < 0.001 **** = 0.00001 < p < 0.0001 Abbreviations: EED = Environmental Enteric Dysfunction; SEEM = Study of Environmental Enteropathy and Malnutrition; AMANHI = Alliance for Maternal and Newborn Health Improvement; MAR04958 = ribose [c] ⇔ ribose [e]; MAR05042 = inosine [c] ⇔ inosine [e]; MAR05071 = lysine [e] ⇒ lysine [c]; MAR09606 = H^+^ [c] + Na^+^ [e] + glycine [e] ⇒ H^+^ [e] + Na^+^ [c] + glycine [c]; MAR06381 = H^+^ [c] + Na^+^ [e] + serine [e] ⇒ H^+^ [e] + Na^+^ [c] + serine [c]; MAR05448 = HCO ^-^ [c] + 2 Na^+^ [c] ⇒ HCO ^-^ [e] + 2 Na^+^ [e]; RIPTiDe = Reaction Inclusion by Parsimony and Transcript Distribution; LDA = linear discriminant analysis; ATP = adenosine triphosphate; NADPH = nicotinamide adenine dinucleotide phosphate; NAD+ = nicotinamide adenine dinucleotide; H^+^ = hydrogen ion; Na^+^ = sodium ion Definitions: Stunted = Length-for-Age Z-Score (LAZ) < -2.0; Not Stunted = Length-for-Age Z-Score (LAZ) ≥ -2.0.

**Supplementary Figure 2.**
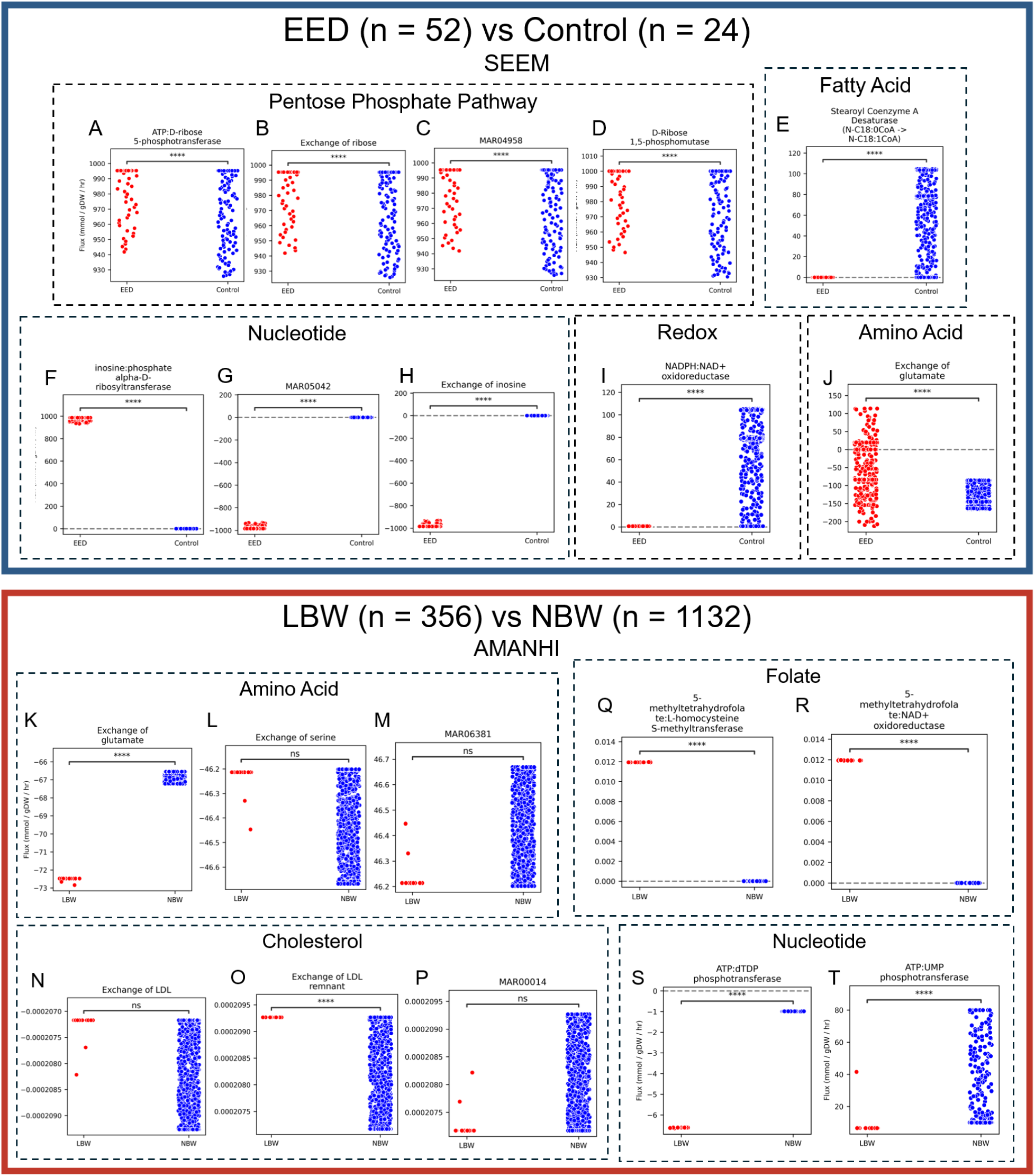
Altered Amino Acid and Nucleotide Metabolism Differentiates EED from Controls and LBW from NBW. Jitter plots showing the top 10 reactions identified by our Reaction Inclusion by Parsimony and Transcript Distribution (RIPTiDe) and Linear Discriminant Analysis framework. Our framework was applied to SEEM duodenal transcriptomics (A-J) and to AMANHI maternal plasma proteomics from visit 2 (24-28 weeks) (K-T). The top 10 reactions were categorized into subsystems by a gastroenterologist based on their expert opinion as to each reaction’s biochemical function. In each plot, the x-axis describes the altered reaction between 1) EED (n = 52) or 2) LBW (n = 356) and 1) controls (n = 24) or 2) NBW (n = 1132). The y-axis shows the flux values sampled from the context-specific metabolic network reconstructions produced by RIPTiDe. A two-sided Mann-Whitney U-test was used to assess the significance of differences in flux distributions. Note that the scale of the y-axes varies between plots and is shown primarily for illustrative purposes. Both sets of discriminating reactions implicate Nucleotide Metabolism (F, G, H, S, T) and Amino Acid Metabolism (J, K, L, M). n.s. = not significant (p >= 0.05) * = 0.01 < p < 0.05 ** = 0.001 < p < 0.01 *** = 0.0001 < p < 0.001 **** = 0.00001 < p < 0.0001 Abbreviations: EED = Environmental Enteric Dysfunction; SEEM = Study of Environmental Enteropathy and Malnutrition; LBW = low birth weight; NBW = normal birth weight; AMANHI = Alliance for Maternal and Newborn Health Improvement; MAR04958 = ribose [c] ⇔ ribose [e]; MAR05042 = inosine [c] ⇔ inosine [e]; MAR06381 = H^+^ [c] + Na^+^ [e] + serine [e] ⇒ H^+^ [e] + Na^+^ [c] + serine [c]; MAR00014 = LDL [e] ⇒ LDL remnant [e] + 680 cholesterol [e] + 1515 cholesterol-ester pool [e]; ATP = adenosine triphosphate; NADPH = nicotinamide adenine dinucleotide phosphate; NAD+ = nicotinamide adenine dinucleotide; LDL = low density lipoprotein; H^+^ = hydrogen ion; Na^+^ = sodium ion Definitions: LBW = birthweight < 2.5 kg; NBW = birthweight ≥ 2.5 kg

**Supplementary Table 1.** Maternal Comorbidities are Not Significantly Associated with Adverse Birth Outcomes or Infant Growth Outcomes at 2 Years. The association between maternal comorbidities (history of chronic hypertension (n = 140), diabetes (n = 19), epilepsy (n = 2), tuberculosis (n = 32), or thyroid disease (n = 12)) with adverse birth outcomes and infant anthropometrics at 2 years of age is presented. Each variable presented is categorical; numerical variables such as birth weight were discretized into clinically meaningful categories prior to statistical analysis. P-values were calculated by a Fisher’s Exact Test with probabilities approximated via Monte Carlo simulation, then adjusted by the Benjamini-Hochberg procedure. After adjustment, no significant (q < 0.05) associations were found. Abbreviations: FDR = false discovery rate

| Variable | FDR q-value |  |  |  |  |
| --- | --- | --- | --- | --- | --- |
|  | Chronic Hypertension<br>(n = 140 yes, n = 2,259 no) | Diabetes<br>(n = 19 yes, n = 2,383 no) | Epilepsy<br>(n = 2 yes, n = 2,400 no) | Tuberculosis<br>(n = 32 yes, n = 2,370 no) | Thyroid Disease<br>(n = 12 yes, n = 2,390 no) |
| **Birth Outcomes and Infant Anthropometrics at Birth** |  |  |  |  |  |
| Pregnancy Outcome | 0.35 | 0.35 | 1.00 | 0.96 | 0.13 |
| Term of Delivery | 0.35 | 0.09 | 1.00 | 0.96 | 0.34 |
| Birthweight | 0.69 | 0.68 | 1.00 | 0.57 | 0.13 |
| Birth Phenotype | 0.35 | 0.35 | 1.00 | 1.00 | 0.47 |
| **Infant Anthropometrics at 24-Month Follow-Up** |  |  |  |  |  |
| Stunting | 0.35 | 0.09 | 1.00 | 0.96 | 0.34 |
| Wasting | 0.35 | 1.00 | 1.00 | 0.57 | 0.62 |

**Supplementary Table 3:** Maternal Metadata is an Insufficient Predictor of Child Growth Outcomes. Five random forest models were trained to predict each of the five proxy outcomes using only maternal metadata (not using proteomics data). The OOB balanced accuracies are presented in the table as an estimate of the models’ accuracies for predicting the outcomes for unseen participants, adjusting for class imbalance. The balanced accuracy score has a minimum value of 0% and a maximum value of 100%, with 50% being the expected accuracy of a blind estimator. The OOB balanced accuracies attained by our random forest models are consistently low – indicative that maternal metadata is an insufficient predictor of any of the proxy outcomes. Abbreviations: OOB = out-of-bag; SGA = small for gestational age; AGA = appropriate for gestational age; LBW = low birth weight; NBW = normal birth weight Definitions: SGA = birthweight < 10^th^ percentile for gestational age; AGA = birthweight ≥ 10^th^ percentile for gestational age; LBW = birthweight < 2.5 kg; NBW = birthweight ≥ 2.5 kg; Stunted at 2 years = length-for-age Z-score < -2.0 at 2 years of age; Not stunted at 2 years = length-for-age Z-score ≥ -2.0 at 2 years of age

| Proxy Outcome | OOB Balanced Accuracy (%) |
| --- | --- |
| SGA (n = 674) vs AGA (n = 1305) | 54.3% |
| Preterm (n = 368) vs Term (n = 2021) | 51.5% |
| LBW (n = 489) vs NBW (n = 1491) | 52.0% |
| Stunted (n = 928) vs Not Stunted (n = 857) at 2 Years | 56.9% |
| Wasted (n = 209) vs Not Wasted (n = 1569) at 2 Years | 50.0% |

## Supplementary Methods

### Sample Collection

#### Cohort A (AMANHI)

Biological material collected from Cohort A participants included maternal blood, urine, and stool samples, placental tissue, and infant cord blood. Trained phlebotomists collected maternal and umbilical cord blood, which was centrifuged to obtain serum, plasma, and buffy-coat aliquots.

Maternal antenatal blood and urine samples were collected at a single timepoint per participant. with women randomized to sample collection at either the 24–28 week or 32–36 week antenatal visit. Placental and umbilical cord tissue were collected at delivery, with a portion flash-frozen and a portion fixed for histological examination. Maternal and newborn stool samples were collected for microbiota analysis. All biospecimens were processed and stored at −80°C following standardized, harmonized protocols implemented across AMANHI sites, and biobanked at the Aga Khan University, Karachi, Pakistan for the Karachi cohort.

#### Cohort B (SEEM)

Duodenal tissue was utilized for bulk RNA sequencing. Due to ethical and logistical constraints precluding obtaining duodenal biopsies from healthy children in this setting, age-matched comparator biopsies were obtained from children undergoing endoscopy at CCHMC for unrelated clinical indications, restricted to those whose biopsy findings and final diagnosis did not support eosinophilic esophagitis, celiac disease, or inflammatory bowel disease, and who had not received antibiotics within four weeks of endoscopy.

### Definition of Growth and Nutritional Status Outcome Groups (EED Proxy Phenotypes)

#### Birth-Defined Outcomes

Three proxy phenotypes were defined using data collected at delivery. 1) Preterm birth, defined as delivery prior to 37 completed weeks of gestation, was compared against term delivery (≥ 37 completed weeks of gestation); 2) Small-for-gestational-age (SGA), defined as a birth weight-for-gestational-age percentile below the 10th percentile, was compared against appropriate-for-gestational-age (AGA) infants (birth weight-for-gestational-age percentile ≥ 10th percentile). 3) Low birth weight (LBW), defined as a birth weight below 2.5 kg regardless of gestational age, was compared against normal birth weight (NBW) infants (birth weight ≥ 2.5 kg).

#### Follow-up-Defined Outcomes

Two proxy phenotypes were defined using anthropometric data collected at the 24-month follow-up visit: 1) Stunting, defined as a height-for-age Z-score (HAZ) below −2.0 relative to the WHO Child Growth Standards, was compared against non-stunted children (HAZ ≥ −2.0). 2) Wasting, defined as a weight-for-height Z-score (WHZ) below −2.0 relative to the WHO Child Growth Standards, was compared against non-wasted children (WHZ ≥ −2.0).

### Biospecimens

#### Cohort A (AMANHI) - Plasma Proteomics

A total of 100 μL of maternal plasma per sample (two aliquots per participant) was shipped from Aga Khan University, Pakistan, to the Max Planck Institute of Biochemistry, Germany, for untargeted proteomic analysis. From each sample, 5 μL of plasma was processed using a perchloric acid workflow with neutralization (PCA-N), which significantly reduces costs with greater proteomic depth. (13). Details of the full proteomics workflow are included in supplemental methods.

The PCA-N workflow was semi-automated using a Bravo liquid handling platform. Plasma (5 μL) was diluted in 25 μL of water, combined with perchloric acid, and incubated in 96-well plates.

Following centrifugation, the supernatant was transferred to a 384-well plate and neutralized to a pH of 8.0–8.5 to provide optimal conditions for enzymatic digestion. Subsequently, proteins were reduced, alkylated, denatured and digested using trypsin.

Following digestion, peptides were loaded onto disposable Evotip C18 trap columns using the Bravo liquid handling platform and separated using an Evosep One liquid chromatography system operated in the 60 samples-per-day (SPD) method. This was coupled to an Orbitrap Astral mass spectrometer (Thermo Fisher Scientific) which employed data-independent acquisition (DIA) technique. Raw DIA data were processed with DIA-NN 1.8.1 against a spectral library predicted in silico with AlphaPeptDeep from the human reference proteome (UniProt UP000005640), and protein-level intensities were obtained with directLFQ and log2-transformed.

#### Cohort B (SEEM) - Duodenal Tissue Transcriptomics

RNA was extracted from duodenal biopsy tissue collected at endoscopy from 52 children with Environmental Enteric Dysfunction (EED) and 24 comparator controls. Sequencing libraries were prepared using polyA RNA selection, followed by fragmentation, complementary DNA (cDNA) synthesis, and adaptor ligation, using the TruSeq RNA Sample Preparation kit (Illumina). Libraries were sequenced as paired-end 75 base pair reads on an Illumina HiSeq 2000 platform at the Cincinnati Children’s Hospital Medical Center (CCHMC) Digestive Health Center Gene Expression Core, with a minimum sequencing depth of 20 million reads per sample (6, 11).

Raw sequencing reads (FASTQ files) were processed using a standardized, cloud-based RNA-sequencing pipeline (37). Reads were aligned to the human reference genome using the

STAR aligner, and gene-level transcript counts were subsequently quantified for use in downstream context-specific metabolic modeling.

### Statistical and Bioinformatic Analysis

#### Generic Genome-scale Metabolic Model (Human1)

Genome-scale Metabolic Models (GEMs) are mathematical models of the human cell which use Gene-Protein-Reaction association rules to link genomic, transcriptomic, or proteomic data to their catalyzed metabolic reactions. GEMs are intentionally generic, able to complete a core list of metabolic functions that any cell should. Several popular GEMs are widespread in literature such as Recon3D (28) and Human1 (29). Our analysis uses Human1 for its rich, open-source community and extensive documentation, although the methods described could theoretically be applied to any similar GEM.

#### Reactome

Reactome (30) is an open-source database of curated, peer-reviewed metabolic pathways. Reactome provides built-in analysis tools for Pathway Enrichment Analysis, determining statistically over-represented pathways from an input list of genes or proteins. We applied Reactome’s analysis tools to over- and under-abundant proteins to identify affected metabolic pathways.

#### Enrichr

Enrichr (31) is a large collection of gene set libraries and is a superset of Reactome. Beyond pathways, Enrichr maps genes to biological processes, diseases, transcription factors, and more. We employed the use of Enrichr libraries to elucidate relevant biological processes related to differentially abundant proteins.

#### Reaction Inclusion by Parsimony and Transcript Distribution (RIPTiDe)

RIPTiDe (32) is an algorithm based on parsimonious Flux Balance Analysis (pFBA) that, given transcriptomic evidence, prunes a large, generic metabolic model to produce a smaller, context-specific metabolic model. RIPTiDe was used for creating context-specific metabolic models based on SEEM transcriptomic or AMANHI proteomic data. Unlike methods that seek maximum consensus with ‘omics data, RIPTiDe first applies an ‘omics-agnostic pFBA approach to maximize the flux through an objective reaction (typically a generic biomass or cell maintenance reaction) and then measures the correlation of the resulting contextualized model to the ‘omics data. This method has been demonstrated to more accurately predict real metabolic activity in *Escherichia Coli* and has been used in similar studies (5, 33), justifying our usage here. Other contextualization algorithms like Cost Optimization Reaction Dependent Assessment (CORDA) (34) or Fast Task-driven Integrative Network Inference for Tissues (ftINIT) (35) are used for proteomics data integration, but require the user to set a known expression threshold a priori or bin proteins in high, medium, and low expression levels (losing much of the granularity that our quantitative proteomics data provides).

#### Linear Discriminant Analysis (LDA)

LDA is a supervised machine learning technique that identifies the linear decision boundary that maximally separates the classes of interest. In our metabolic modeling analysis, this technique was chosen to identify the most important reaction-level differences in context-specific metabolic models.

#### Data Pre-Processing

Plasma proteomic samples missing values for more than 40% of proteins, and proteins missing values in more than 10% of samples, were excluded from analysis. Remaining missing values were imputed using Iterative Singular Value Decomposition (SVD) (36) with a rank empirically determined to minimize a bootstrapped estimate of the matrix reconstruction error. To confirm that imputation did not introduce statistical bias, the distributions of each protein’s abundance values were compared before and after imputation using a two-sample Kolmogorov-Smirnov test, with a Bonferroni correction applied at a significance threshold of 0.05. Following imputation and post-hoc validation, plasma samples from the third antenatal visit (32–36 weeks gestation) were excluded from downstream analysis due to low sample size. To strengthen downstream comparisons, analysis was further restricted to participants with plasma samples available at both the first (8–19 weeks) and second (24–28 weeks) antenatal visits.

#### Differential Expression Analysis

Protein-level differences between maternal plasma proteomic subcohorts were identified using Mann-Whitney U tests with a Benjamini-Hochberg correction, at a significance threshold of 0.05. A protein was considered differentially abundant between two subcohorts if its abundance distributions differed significantly and the absolute log2 fold change was ≥ 1.0.

#### Pathway Enrichment and Over-Representation Analysis

To infer biological processes underlying differential protein abundance, over-representation analysis (ORA) was performed using Reactome and Enrichr. Sets of significantly upregulated and downregulated proteins were analyzed against Reactome pathways and Enrichr libraries to identify enriched molecular processes. If no such proteins existed, the log2 fold change was relaxed from 1.0 to ≈ 0.14 (equivalent to a 10% fold change) to permit analysis with Reactome and Enrichr. Enrichment significance was assessed using multiple testing-adjusted p-values (FDR). This enabled orthogonal interrogation of dysregulated pathways, strengthening confidence in pathway-level inference.

### Random Forest Modeling

Random forests are a popular machine learning method that extend decision trees to reduce variance and provide robust predictions. To measure the value of maternal metadata in predicting growth proxies, we trained random forest models on maternal metadata (without proteomics) and measured the Out-Of-Bag (OOB) balanced accuracies as a performance metric.

### Sensitivity Analysis

A subcohort of the AMANHI participants who had at least one comorbidity (chronic hypertension, diabetes, epilepsy, tuberculosis, thyroid disease) were investigated for their potential impact on the larger cohort. Fisher’s Exact Tests with 10 million rollouts each and a Benjamini-Hochberg correction were applied to identify significant (FDR < 0.05) associations between these comorbidities and our five growth proxies.

We also conducted a sensitivity analysis by excluding these participants from the larger AMANHI cohort and repeating the metabolic modeling methods described above. The reactions identified from the sensitivity analysis were compared with reactions identified using the entire cohort to study the influence of these participants on the overall pipeline and characterize the pipeline’s sensitivity to anomalies.

### Ethical Approval and Consent

SEEM was approved by the Aga Khan University Ethical Review Committee (ERC: 2446 Ped ERC-13) and the Institutional Review Board at Cincinnati Children’s Hospital Medical Center (CCHMC, Study ID 2016-0387), and is registered with ClinicalTrials.gov (NCT03588013).

AMANHI was approved by Aga Khan University Ethical Review Committee as well as by the World Health Organization Ethics Review Committee (RPC 532). Written informed consent was obtained from all participants (or their parent/legal guardian, for pediatric participants) prior to enrollment in each study.

### Data and Code Availability

All figures were generated and analysis was performed in Python 3.14.3 using libraries including conda (26.7.2), pip (26.0.1), numpy (2.4.6), pandas (2.3.3), cobra (0.31.1), fisher-rxc (1.0.2), matplotlib (3.10.9), riptide (3.4.81), seaborn (0.13.2), statannotations (0.7.2), gurobi (13.0.2), scikit-learn (1.8.0), umap-learn (0.5.12), scipy (1.17.1), fancyimpute (0.7.0), and tqdm (4.67.3). Additional information required to re-analyze the reported data is available from the lead author on request.

The code written to conduct the analysis is available at https://github.com/SyedLab-GI/CompEE Transcriptomic data for the SEEM study is available at GEO Accession code GSE159495.

Proteomic data for the AMANHI study is not publicly available, but may be requested from the lead author (FJ, and IN).

### Supplementary Results

#### Proteomics Data Pre-Processing

Iterative SVD with a rank of 66 was found to minimize a bootstrapped estimate of the matrix reconstruction error for imputing missing proteomics values. The distribution of five proteins were found to be significantly (FWER < 0.05) affected by a two-sample Kolmogorov-Smirnov test. However, upon investigation, the effect sizes were all negligible (between 0.05 and 0.06, allaying concerns of bias from imputation.

2,458 plasma samples were collected at visit 1, 1,890 at visit 2, and 235 were collected at visit 3. Only the 1,864 participants who had plasma samples collected at both visits 1 and 2 were retained for differential expression, pathway enrichment, and metabolic modeling analyses.

#### Differentially Abundant Proteins Across EED Proxy Comparisons

Differential Expression Analysis revealed statistically significant differences in protein abundance between SGA and AGA, LBW and NBW, Stunted at 2 Years and Not Stunted at 2 Years, and Wasted at 2 Years and Not Wasted at 2 Years cohorts. However, none of these differences met the log2 fold change threshold of 1.0 for biological significance. No significant differences (either statistically or biologically) were found for the Preterm and Term cohort.

#### Pathway Enrichment Analysis

Since no proteins met the log2 fold change threshold, the threshold was relaxed to ≈ 0.14 (equivalent to a 10% fold change) for downstream pathway enrichment analysis. Exploratory ORA conducted via Reactome and Enrichr identified up-regulation of immune activation and tissue remodeling and down-regulation of barrier integrity and cellular homeostasis in mothers whose children were SGA, up-regulation of Extracellular Matrix (ECM) remodeling in the LBW cohort, up-regulated inflammatory response and decreased lymphocyte migration in the stunting cohort, and increased cell adhesion in the wasting cohort.

#### Predicting Growth Proxies from Maternal Metadata

Across all five proxies, the Random Forest models exhibited near coin-flip accuracy, indicating that, without proteomics data integration, maternal metadata alone was an insufficient predictor of these growth proxies **(Supplementary Table 2).**

#### Sensitivity Analysis

7.8% of mothers in the AMANHI cohort (n = 194) had a history of comorbidities known to adversely affect pregnancy (chronic hypertension (n = 140), diabetes (n = 19), epilepsy (n = 2), tuberculosis (n = 32), or thyroid disease (n = 12)). Categorical associations between presence of each comorbidity and each of the five proxies as well as pregnancy outcome (live birth, stillbirth, or miscarriage) were tested. No significant (FDR < 0.05) associations were found **(Supplementary Table 1)**.

This subcohort was further interrogated using the differential expression, pathway enrichment, and metabolic modeling pipelines utilized for the five proxies of EED described earlier.

Differential expression again revealed statistically – but not biologically – significant differences in protein abundance in mothers with one or more of these comorbidities. After relaxing the log2 fold change threshold to ≈ 0.14, analysis of upregulated and downregulated proteins revealed affected immune system activity, hemostasis, metabolism of carbohydrates and carbohydrate derivatives, and metabolism of water-soluble vitamins. Metabolic modeling with RIPTiDe and Human1 primarily implicated Vitamin A metabolism and Amino Acid Metabolism in the top 10 identified reactions, corroborating the pathway enrichment analysis findings.

To identify the impact of these participants on the larger cohort, a sensitivity analysis was conducted wherein the differential expression, pathway enrichment, and metabolic modeling analyses were repeated with these 194 participants removed. Although the specific reactions identified by LDA, the subsystems associated with the top 10 reactions remained highly similar, validating the pipeline’s resilience against outliers.

## Notes

https://github.com/SyedLab-GI/CompEE

https://www.ncbi.nlm.nih.gov/geo/query/acc.cgi?acc=GSE159495

### Competing Interest Statement

The authors have declared no competing interest.

